# A fluorescence-based sensor screen identifies MED12 as a potential microsatellite instability regulator in colon cancer

**DOI:** 10.1101/2023.08.17.553681

**Authors:** João M. Fernandes Neto, Subramanian Venkatesan, Matheus Dias, Cor Lieftink, Ben Morris, Kaspar Bresser, Loredana Vecchione, Bastiaan Evers, Ferenc Scheeren, Ton Schumacher, Roderick L. Beijersbergen, René Bernards

## Abstract

Inactivation of the DNA mismatch repair (MMR) system, due to (epi)genetic alterations of MMR genes, increases the frequency of mutations across the genome, creating a phenotype known as microsatellite instability (MSI). Cancers with this phenotype have been associated with a better prognosis for some time, but only since recently it has been recognised as a predictive biomarker of response to immunotherapy. Because MSI tumours accumulate more insertions and/or deletions in coding regions of the genome containing microsatellites, there is an increase in neoantigens resulting from reading frame shifts, which promotes immunogenicity. To investigate if additional genes exist that can cause an MSI phenotype, we developed a fluorescence-based sensor to identify genes whose inactivation increases the rate of frameshift mutations on microsatellite sequences in cancer cells. Using genome-scale CRISPR/Cas9 screens, we identified *MED12* as a potential new regulator of microsatellite instability. Consistent with this, we found that *MED12* mutant colon cancers that lack mutations in the known MMR genes are more likely to be of the MSI phenotype.

## Introduction

Microsatellites are repetitive DNA tracts in which certain DNA motifs (ranging from one to several base pairs in length) are repeated, typically 5 to 50 times (1,2). Microsatellites are abundant in the genome; however, they occur mainly in non-coding DNA. Replication slippage, caused by the transient dissociation of the replicating DNA strands followed by misaligned re-association, occurs frequently in microsatellites but mutations are generally corrected by the mismatch-repair (MMR) system. However, in the absence of a proficient MMR system, due to genetic or epigenetic alterations of any of the four well established MMR genes (*MSH2, MSH6, MLH1* and *PMS2*), the rate of mutations will significantly increase, leading to a phenotype known as microsatellite instability (MSI) (3). MMR deficiency, and therefore microsatellite instability, increases the chances of developing cancer (4), but on the other hand, some patients with MSI-H tumours, like early-stage colorectal cancer patients, have a better prognosis compared to microsatellite stable (MSS) patients (5). With the recent advances in immunotherapy, the life expectancy of MSI patients has improved even further, as MSI tumours have been shown to respond better to immune checkpoint blockade therapies as compared to conventional chemotherapeutics (6). Because loss of MMR increases the mutation rate in tumours, the rate of putative frameshift peptide neoantigens also increases. Frameshift mutations are genetic mutations caused by insertions or deletions of several nucleotides in a DNA sequence that is not divisible by three (7). Due to the triplet nature of gene expression by codons, such a mutation can result in altered transcript and peptide stretches, which can lead to a more immunogenic tumour microenvironment (8,9).

It has been almost 10 years since the so-called MSI-like phenotype was identified in colorectal cancer (10). Tumours with this phenotype score negative for MSI in a clinical diagnostic assay, but have an expression profile similar to MSI tumours and, like MSI tumours, have a better prognosis and exhibit an increased lymphocytic infiltrate. More recently, a study also showed that a fraction of MSS tumours has a high immunoscore and better prognosis (11). Together, these data raise the possibility that there might be additional (currently unknown) genes involved in the regulation of DNA mismatch repair. Because in CRC a positive MSI test or dMMR is the eligibility criteria for immunotherapy administration, some patients who could benefit from immunotherapy might currently not be identified.

We developed an MSS colon cancer cell line harbouring a fluorescent-reporter that becomes irreversibly activated when a slippage event occurs within a microsatellite sequence in the reporter. We used this cell line to perform a whole-genome CRISPR/Cas9 screen to identify novel genes that regulate MMR in order to increase the number of biomarkers that we can use in the clinic to decide which patients should receive immunotherapy and to, possibly, explain the MSI-like phenotype that is observed in some colon cancer patients.

## Results

Microsatellites, because of their repetitive nature, are more prone to frameshift mutations than other genomic regions (12). To detect replication slippage within microsatellites we generated a frameshift-mutation sensor plasmid. This plasmid (Figure 1A) consists of a so-called “MSI tract” – a DNA sequence consisting of 23 guanine repeats (G23) – downstream of a constitutive promoter, followed by the Cre recombinase gene. We designed the MSI tract such that the Cre recombinase gene would be out-of-frame. This way, Cre protein expression is linked to the occurrence of in-frame mutants through replication slippage. We named this vector the G23 MSI activator. Because of the high mutation rate in microsatellites, it is possible that after being mutated in-frame, a subsequent mutation could place the Cre gene again out-of-frame. To ensure that a one-time activation of the MSI activator would lead to an irreversible mark in the cell, we generated a second vector – called MSI reporter. Here, a selection marker (neomycin) and a red fluorescent protein (katushka) were cloned in-frame, after a floxed non-sense ORF. With this double system, upon in-frame mutation of the MSI activator, Cre expression leads to excision of the non-sense ORF, making the cells both irreversibly resistant to neomycin and katushka-positive (Fig. 1A). We also included selection markers in these reporters (hygromycin and blasticidin, respectively) to facilitate selection of cells with the double integration.

**Figure 1:**
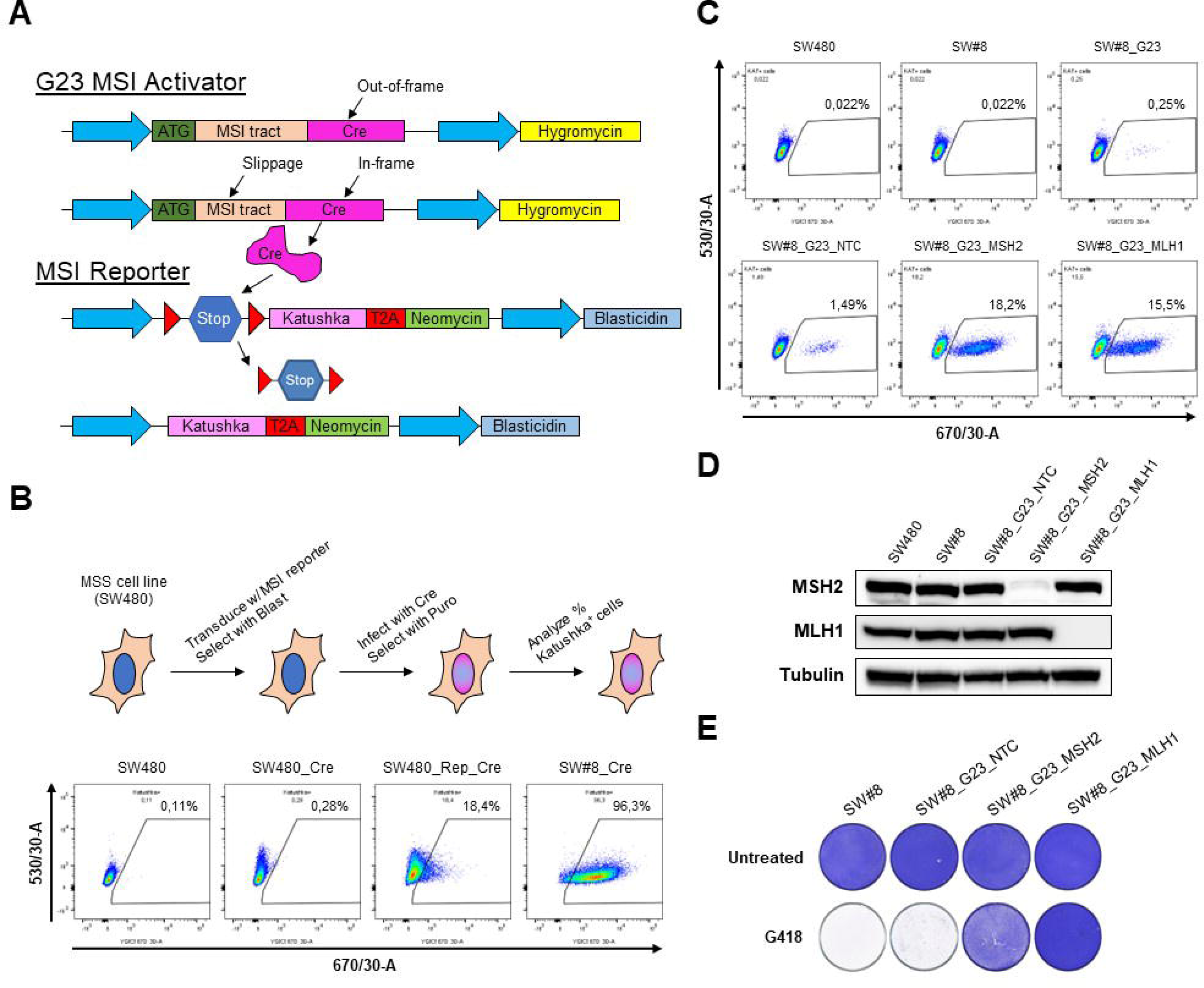
Development of the fluorescent-based sensor to study microsatellite instability. A. Schematic representation of the MSI sensor. B-E. Validation of the MSI sensor in MSS CRC cell line SW480. In (B) SW480 cells were transduced with the MSI reporter and several cloned were generated; the clones and the polyclonal population were then transduced with a plasmid expressing Cre recombinase (Addgene #30205). Activation of the reporter upon constitutive Cre expression was measured by flow cytometry; the best clone (SW#8) was selected for further studies. In (C-E) SW#8 cells were transduced with the G23 MSI activator and with sgRNAs targeting positive control genes (MSH2 and MLH1) or a non-targeting control (NTC). In (C) activation of the reporter was measured by flow cytometry after 3 weeks in culture. Y axis = Blue 530/30-A; X axis = Yellow-green 670/30-A; gates indicate katushka-positive cells; percentages of katushka-positive cells are highlighted above the gates. In (D) protein was isolated from the cell lines, as indicated, to assess levels of MSH2 and MLH1 by western blot. Tubulin was used as a loading control. A representative image from two biological replicates is displayed. In (E) G418 (200µg/mL) was added to the indicated cells. 10 days later plates were fixed and stained.

We used the MSS colorectal cancer cell line SW480 (13) to validate our MSI sensor system. First, we transduced SW480 cells with the MSI reporter; through single-cell sorting we were able to pick a clone (called SW#8) in which the sensor became uniformly activated upon Cre induction (Fig. 1B). Next, we transduced the SW#8 cells with the G23 MSI activator, generating SW#8_G23. As a positive control, we knocked-out two established MMR genes in these cells to test if this would result in significant activation of the sensor. We observed that the knockout (KO) of MLH1 or MSH2 significantly increased the activation of the MSI sensor over time, compared to the control (Figs. 1C-D). Additionally, all the katushka-positive cells were resistant to G418 (Fig. 1E). However, it is important to note that it took over 3 weeks to see a 10-fold induction relative to the control. This is not unexpected, as a frameshift mutation needs to occur within the 23 base pair region of the “MSI tract” out of the many other microsatellites in the genome, to be activated. Therefore, the more DNA replication cycles the cells undergo, the higher the chances of activating the sensor. The estimated slippage mutation rate for this type of microsatellite is 1E-4 (14).

To find potential new regulators of microsatellite instability, we performed a genome-wide CRISPR/Cas9 screen in SW#8_G23 cells. We hypothesised that regulators of MSI would deregulate the MMR system. Because MMR-deficient cells are resistant to temozolomide (15) (Sup. Fig. 1A), we also included an arm in the screen where the cells were treated with temozolomide (TMZ) to enrich for MMR deficient cells. We generated Cas9 expressing SW#8_G23 cells and transduced them with the Brunello sgRNA library. Because in MMR-proficient cells slippage on any single microsatellite is a rare event, we kept the cells in culture for 36 days to increase the chance of activating the sensor in enough cells through the acquisition of an MSI phenotype as a consequence of a gene knockout. We then selected the cells which had activated the sensor with G418 for 12 days (Fig. 2A). In the TMZ arm we also harvested cells before selection with G418, to rule out any side-effect from G418. After this, cells were harvested and the barcodes associated with the gRNAs were recovered by PCR and quantified by deep sequencing, as described earlier (16). The screen performed well technically, as judged by the depletion of essential genes over non-essential genes (Sup. Fig. 1B). As expected, treating the cells with TMZ enriched for sgRNAs targeting MMR genes (Fig. 2B). Not surprisingly, these genes were also enriched in the cells that were harvested after G418 selection, in which the sensor was activated (Sup. Fig. 1C and Table S1). However, in the unbiased arm (untreated), only PMS2 scored as a hit (Fig. 2C). The other MMR genes scored within the top 500, but below the significance threshold of FDR < 0.1 (Table S2). This suggests that in the absence of a strong selection pressure for an MSI phenotype (as was used in the TMZ resistance arm of the screen) the slippage events on the MSI tract of the MSI activator plasmid are too infrequent to yield a signal in a genome-scale genetic screen.

**Figure 2:**
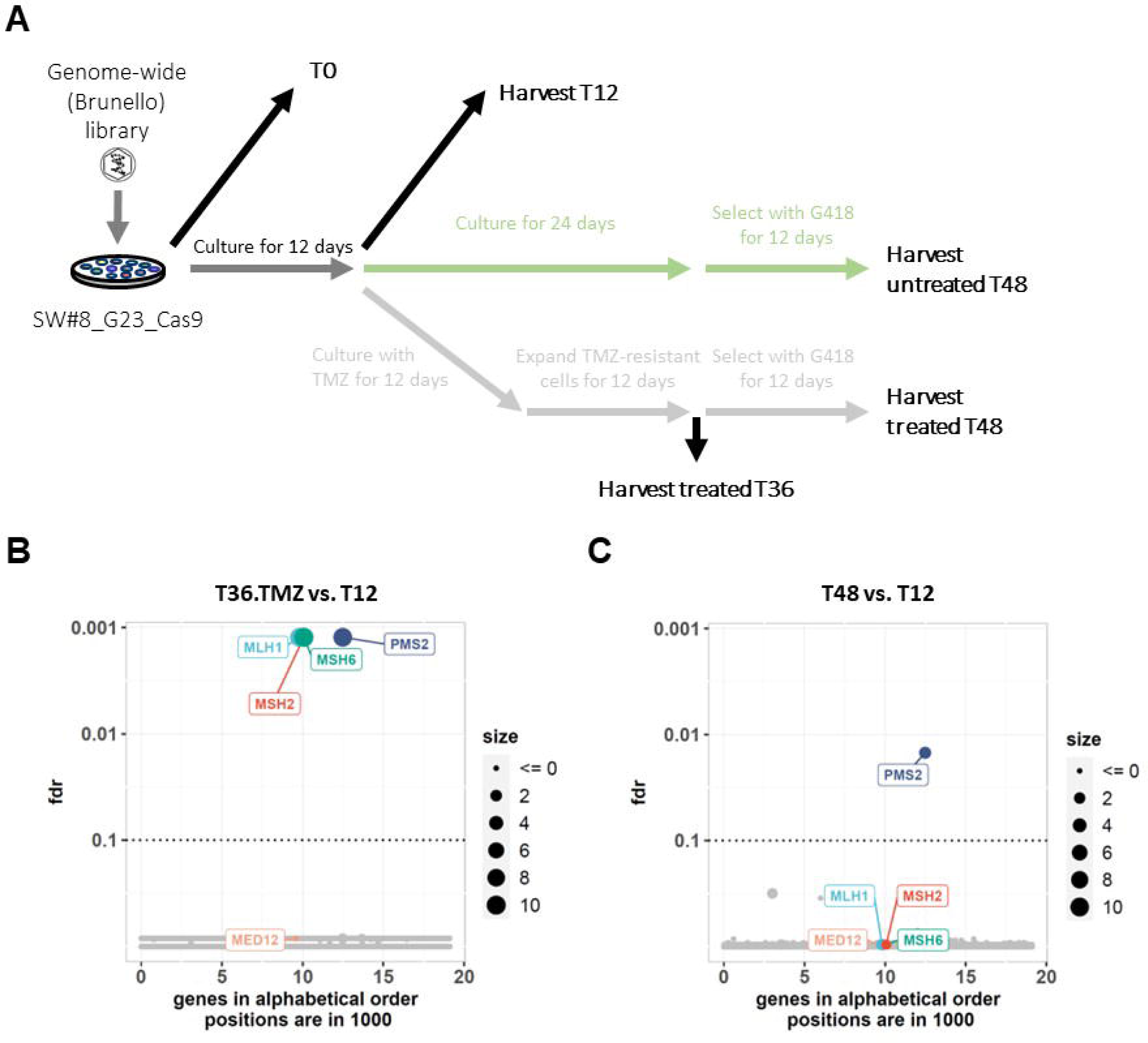
**Genome-wide, fluorescent-based sensor screen is not able to pick regulators of MSI**. A. Schematic representation of the genome-wide CRISPR screen. B, C. Robust rank analysis of the sgRNAs enrichment in the genome-wide screen. Cas9 expressing SW#8 cells were screened with the Brunello whole-genome sgRNA library. In (B) Robust rank results for enrichment of each gene in the Temozolomide treated arm (T36) compared to the reference (T12) is displayed. In (C) Robust rank results for enrichment of each gene in the untreated arm (T48) compared to the reference (T12) is displayed.

To improve the signal-to-noise ratio, we decided to perform a second screen in the same cells using a smaller “MSI-focused” library comprised of 3089 sgRNAs (versus 77441 sgRNAs present in the Brunello library). The new library targeted 528 “MSI-focused” genes (the top enriched genes from our whole genome screens, as well as candidate genes from literature), each with 5 sgRNAs, plus control genes. (Table S3). We transduced the Cas9 expressing SW#8_G23 cells with the MSI-focused gRNA library. As a control, we harvested cells at day 7, to assess the depletion of essential genes over non-essential genes. To increase the chances of activating the MSI sensor, cells were cultured for either 44 or 85 days (Fig. 3A). As early as day 44, all 4 established MMR genes scored as hits in the screen (Fig. 3B). Importantly, one additional gene, not previously connected to MMR, scored as a hit in the screen – MED12 (Fig. 3B). Culturing the cells for longer than 44 days did not yield any additional hits. However, at the later time point, MSH6 didn’t reach the significance threshold of FDR < 0.1 (Fig. 3C and Table S4).

**Figure 3:**
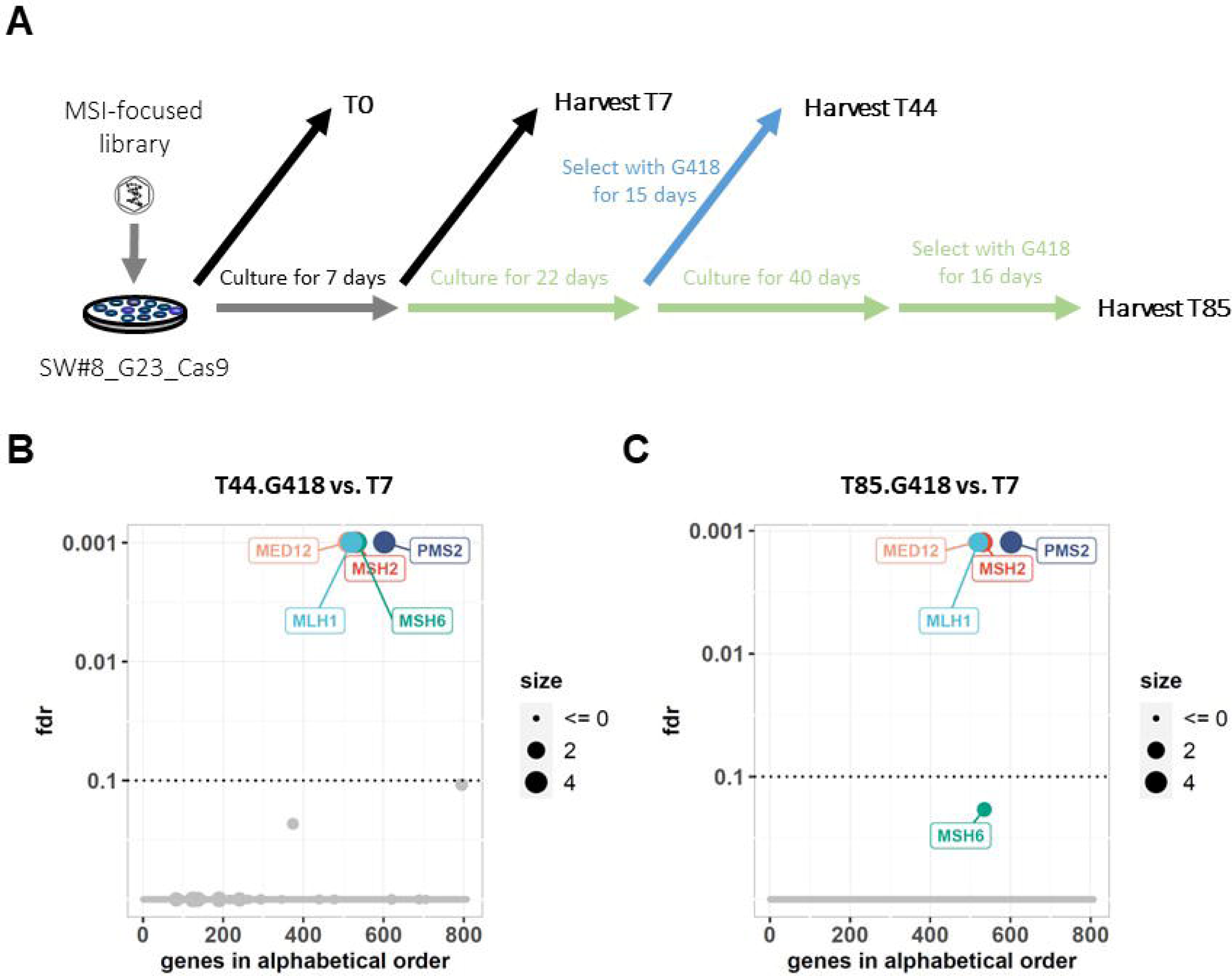
**Focused sgRNA library, fluorescent-based sensor screen picks MED12 a novel regulator of MSI**. A. Schematic representation of the MSI-focused CRISPR screen. B, C. Robust rank results for enrichment of each gene in the MSI-focused screen. Cas9 expressing SW#8 cells were screened with the MSI-focused sgRNA library. The Robust rank distribution of the enriched sgRNAs after 44 and 85 days in culture (B and C, respectively) compared to the reference (T7) is displayed.

MED12 is a component of the MEDIATOR complex, which links transcription factors to RNA polymerase II. Mutations in MED12 are frequent in uterine leiomyomas (17). To further study whether MED12 regulates microsatellite instability, we generated MED12 KOs in SW#8_G23 cells, as well as KOs of MSH2 and MLH1 as positive controls (Fig. 4A). We observed that, on average, loss of MED12 led to a 4-fold increase in MSI sensor activation after 3 months in culture, as compared to control cells. However, far less than the 145-fold increase observed in the positive controls (Fig. 4B). We also observed that KO of MED12 leads to resistance to temozolomide relative to control cells, albeit to a lesser extent than knockout of MLH1 or MSH2 (Fig. 4C). We also tested the MSI status of the MED12 clones using the Promega MSI Analysis System, which is the gold standard MSI assay in clinical research (18). Using this PCR-based method, we tested five nearly monomorphic mononucleotide repeat markers (BAT-25, BAT-26, MONO-27, NR-21 and NR-24). By evaluating the length of these markers, it is possible to detect contractions and expansions. Scoring positive for at least 2 markers is the criteria for classifying a sample as MSI-high (MSI-H) and one marker positive as MSI-low (MSI-L). The positive controls, MLH1 and MSH2, scored positive for 2 markers (BAT-25 and Mono-27), whereas one of the four MED12 KO clones scored positive for 2 markers and one for one marker (Fig. 4D). Thus, by the Bethesda guidelines (19), one MED12 KO line scores as MSI-L, one as MSI-H and two as MSS.

**Figure 4:**
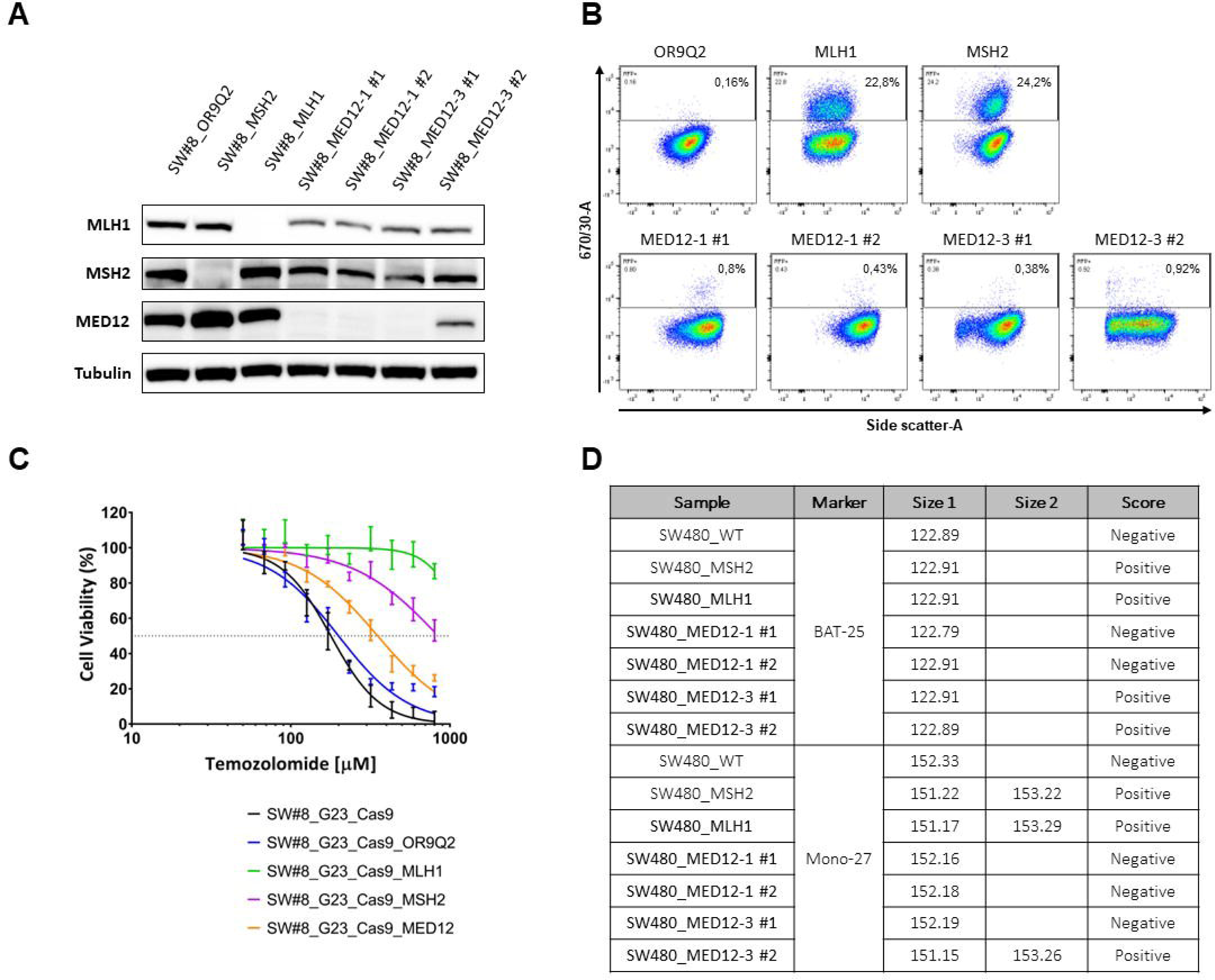
**Validation of MED12 as a potential regulator of microsatellite instability**. A-D. SW#8_G23_Cas9 cells were transduced with sgRNAs targeting positive control genes (MSH2 and MLH1), a negative control gene (OR9Q2) and 2 different sgRNAs targeting MED12. For each MED12 sgRNA 2 different clones were generated (#1 and #2). In (A) protein was isolated from the cell lines, as indicated, to assess levels of MSH2, MLH1 and MED12 by western blot. Tubulin was used as a loading control. A representative image from three biological replicates is displayed. In (B) activation of the reporter was measured by flow cytometry after 3 months in culture. Y axis = Yellow-green 670/30-A; X axis = Side scatter-A; gates indicate katushka-positive cells; percentages of katushka-positive cells are highlighted in the gates. In (C) cells were cultured with increasing concentrations of Temozolomide for 4 days, after which cell viability was measured using CellTiter-Blue®. Standard deviation (SD) from 3 biologically independent replicates (each with 3 technical replicates) is plotted. In (D) the MSI status of the cells was tested by PCR. The results for the BAT-25 and Mono-27 markers are displayed.

As MED12 is primarily known for its role in regulation of transcription, we tested if MED12 knockout affected the expression of the known MRR genes in these cells. We observed that MED12 KO cells slightly downregulated MLH1 and MSH2 expression in SW480 cells (Fig. 4A).

To study the possible relevance of our findings for human colon cancer, we studied the TCGA colon adenocarcinoma cohort. We investigated whether there is a correlation between MED12 mutations and MSI status in these tumours (Fig. 5A-D). This analysis showed that MED12 mutant tumours that lacked other MMR mutations are significantly more likely to be MSI-high relative to MED12 wild type tumours (Fisher’s exact test, P = 0.018) (Fig. 5E). Moreover, we found that the SBS6 mutation signature, which is characteristic of MSI-H tumours (20), is also enriched in MED12 mutant tumours (Fig. 4F). Finally, consistent with the presence of an inflammatory gene signature in MSI-H tumours, we also see that MED12 mutant colon adenocarcinomas have an increased expression of inflammatory gene signatures (Sup. Fig. 2).

**Figure 5:**
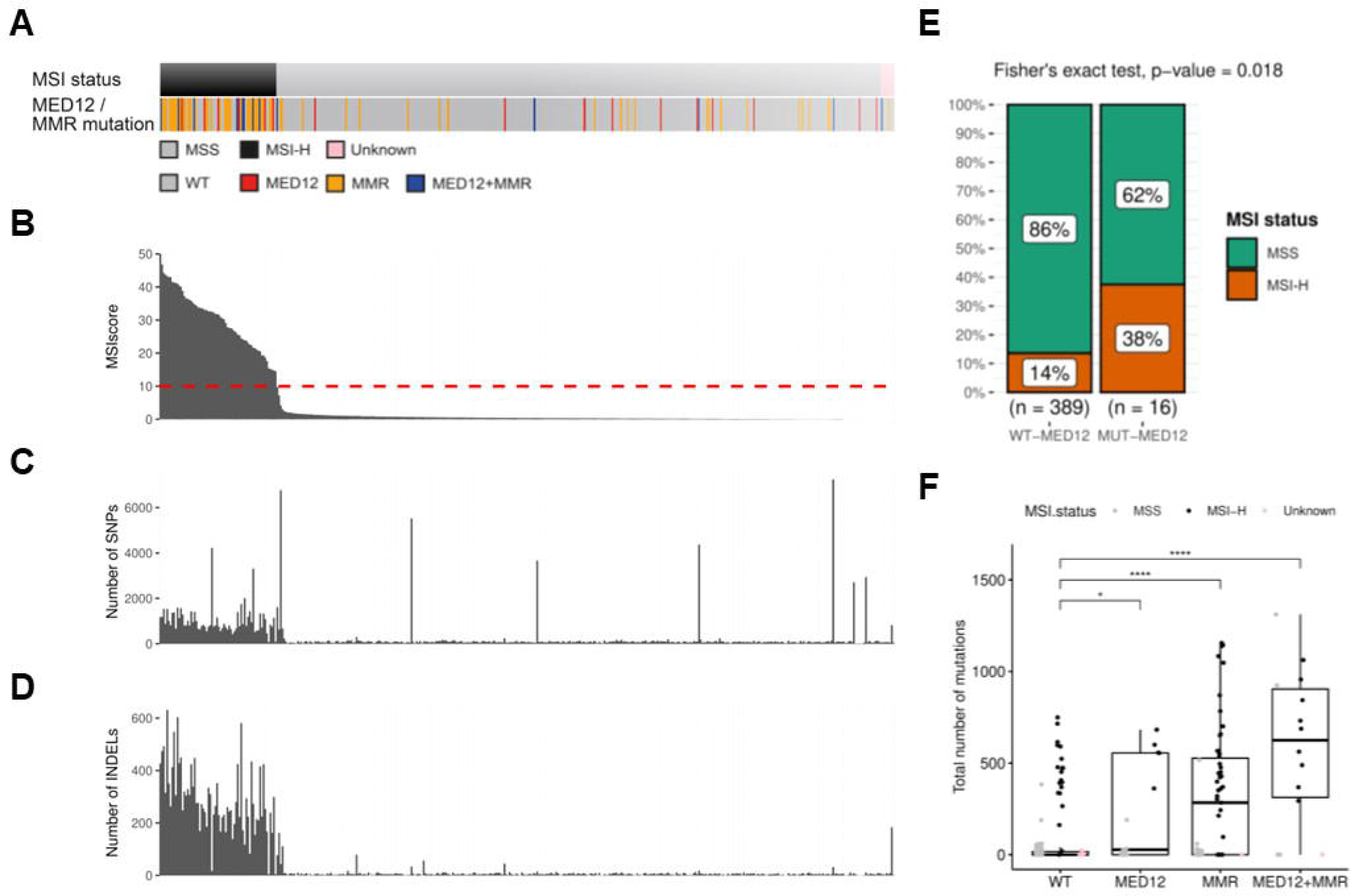
**MED12 mutant colon adenocarcinomas associate with MSI features**. A-D. The plots are aligned and sorted by tumour samples according to their MSI score. In (A) annotation bars of the MSI status (top) and MED12 and MMR gene mutation status (bottom). In (B) MSI score of the TCGA colon adenocarcinoma cohort. Plot is sorted by MSI score. In (C) number of non-synonymous SNPs per tumour sample. In (D) number of INDELs per tumour sample. E, MED12 mutant tumours are significantly enriched for an MSI-high status relative to MED12 wild type tumours (Fisher’s exact test, P = 0.018). MED12 mutant tumours which also had MMR mutations were excluded from this analysis. F, Total number of SBS6 mutations per tumour according to their MED12 and MMR gene mutation status. The colour of the dot indicates the MSI status. MED12 mutant tumours (excluding samples with mutations in the well-established MMR genes, so “MED12 mutant only”), MMR gene mutant tumours and MED12+MMR gene double mutant tumours are significantly enriched with SBS6 mutations (2-sample Wilcoxon test, *FDR ≤ 0.05, ****FDR ≤ 0.0001).

## Discussion

In this study we aimed to identify novel regulators of microsatellite instability using a fluorescent-based sensor and CRISPR screening approach. Our findings suggest that MED12 might play a role in microsatellite instability by downregulating members of the MMR system. MED12 is a component of the mediator of RNA polymerase II transcription (MEDIATOR) complex. As an essential component of the RNA polymerase II general transcriptional machinery, it plays a crucial part in the activation and repression of transcription initiation (21,22). This can potentially explain how MED12 could be involved with microsatellite instability regulation, i.e., MED12 loss might impair the transcription of the MMR genes, causing expression downregulation. However, this downregulation is only “mild”, potentially explaining why the MSI phenotype takes longer to appear, in line with the differences observed in the activation of the MSI sensor in the MED12 KO cells compared to the KO of MSH2 or MLH1. Additionally, EP300, which is also a regulator of MLH1 expression (23) was also in the top 8 genes which were enriched in the MSI-focused screen (Table S4). The link between MED12 and MMR raises the question whether MED12 could indeed contribute to microsatellite instability in CRC tumours. Bioinformatics analysis show MED12 mutant colon adenocarcinomas are enriched for a MSI phenotype in colon adenocarcinoma patients, but more work is necessary to better understand how MED12 loss can lead to microsatellite instability. It should be noted in this context that in our artificial cell model in vitro, the cells go through relatively few DNA replication cycles compared to emerging cancers. Since we observe a small, but measurable effect of MED12 loss even in short term cell culture experiments, it is well possible that effects of MED12 loss in emerging cancers on MSI phenotype is much more pronounced.

Lynch syndrome is a hereditary condition caused by germline inactivation in one of the MMR genes. This condition increases the chances of developing cancer, and because of the accumulation of multiple mutations over time patients also develop MSI tumours. Another condition that causes MSI tumours is called Lynch-like syndrome. Cancers from Lynch-like syndrome patients show MSI but the mechanism for the generation of MSI is unknown, because they have no germline mutations in the MMR genes (24). In a clinical study which analysed tumours from patients with Lynch-like syndrome, *MED12* was found to be mutated in 29% of the tumours (25), which is significantly higher than the 5% mutation frequency observed in CRC patients (26), providing some clinical evidence linking MED12 and MSI. Similar to Lynch-like tumours, MSI-like tumours display a MSI signature but have no germline mutations in the DNA MMR genes (10). This is in line with the finding from our study in which only one clone scored as MSI positive, but all clones downregulated the MMR genes. As a transcriptional regulator, it is plausible that alterations in MED12 could cause this.

Recently, inactivation of MMR genes was shown to trigger dynamic neoantigen evolution and increased response to immunotherapy (27). Interestingly, we observed that MED12 mutant colon adenocarcinoma patients display an increased expression of inflammatory gene signatures, in line with what is observed in immunogenic tumours. In future work, it would be interesting to study whether loss of MED12 contributes to an increase in immunogenicity in vivo. This may be particularly relevant for uterine leiomyoma, as these tumours have a high *MED12* mutation frequency (17).

In the clinic, MSI status can be assessed by 3 methods: immunohistochemistry (IHC) for the MMR proteins MLH1, MSH2, MSH6 and PMS2, (PCR)-based assessment of microsatellite alterations using five microsatellite markers including at least BAT-25 and BAT-26, and next-generation sequencing (28). The first two are significantly cheaper, therefore it is not surprising that in most hospitals the latter is not performed. However, there is increasing evidence suggesting that a subset of patients which score negative for dMMR and MSI could also benefit from immunotherapy. Due to economic reasons, it is not possible to perform next-generation sequencing on every patient that comes into the clinic to identify such a patient subset. In our study we tried to identify markers which could help identify this subset of patients without the need for next generation sequencing approaches.

In summary, we developed a new system to study the development of microsatellite instability. Using this system, together with CRISPR screening technology, we could identify MED12 as a potential new MSI regulator. Since in most hospitals MSI testing is done by immunohistochemistry our findings indicate that it could be relevant to also assess the expression of MED12 by immunohistochemistry in tumours which score negative for dMMR and MSI. In case patients have low expression of MED12 they should then be further evaluated. This would give more patients the opportunity to receive the best possible treatment. Finally, we note that the screening system we developed could also be useful to identify small molecules that can induce an MSI phenotype. That some drugs can induce an MSI phenotype in MSS tumours was recently demonstrated (29). Such small molecule inducers of an MSI phenotype are candidates to sensitize to checkpoint immunotherapy (30).

## Materials and methods

### Cell culture and drug response assays

SW480 cells were cultured in RPMI medium (Gibco 21875034). All the cell lines media were supplemented with 10% FBS (Serana), 1% penicillin/streptomycin (Gibco 15140122) and 2_JmM L-glutamine (Gibco 25030024). All cell lines were cultured at 37°C and with 5% CO2. All cell lines were validated by STR profiling and mycoplasma tests were performed every 2-3 months.

All drug-response assays were performed in triplicate, using black-walled 384-well plates (Greiner 781091). Cells were plated at the optimal seeding density and incubated for approximately 24 hours to allow attachment to the plate. Drugs were then added to the plates using the Tecan D300e digital dispenser. 10 µM phenylarsine oxide was used as positive control (0% cell viability) and DMSO was used as negative control (100% cell viability). Four days later, culture medium was removed and CellTiter-Blue (Promega G8081) was added to the plates. After 1-4 hours incubation, measurements were performed according to manufacturer’s instructions using the EnVision (Perkin Elmer).

### Western Blots

After the indicated culture period, cells were washed with chilled PBS and then lysed with RIPA buffer (25mM Tris – HCl pH 7.6, 150mM NaCl, 1% NP-40, 1% sodium deoxycholate, 0.1% SDS) containing protease inhibitors (Complete (Roche) and phosphatase inhibitor cocktails II and III). Samples were then centrifuged for 10 minutes at 14.000 rpm at 4°C and supernatant was collected. Protein concentration of the samples was normalized after performing a Bicinchoninic Acid (BCA) assay (Pierce BCA, Thermo Scientific), according to the manufacturer’s instructions. Protein samples (denatured with DTT followed by 5 minutes heating at 95°C) were then loaded in a 4-12% polyacrylamide gel. Gels were run (SDS-PAGE) for approximately 60 minutes at 165 volts. Proteins were then transferred from the gel to a polyvinylidene fluoride (PVDF) membrane, using 330 mA for 90 minutes. After the transfer, membranes were placed in blocking solution (5% bovine serum albumin (BSA) in PBS with 0,1% Tween-20 (PBS-T). Subsequently, membranes were probed with primary antibody in blocking solution (1:1000) and left shaking overnight at 4°C. Membranes were then washed 3 times for 10 minutes with PBS-T, followed by one hour incubation at room temperature with the secondary antibody (HRP conjugated, 1:10000) in blocking solution. Membranes were again washed 3 times for 10 minutes in PBS-T. Finally, a chemiluminescence substrate (ECL, Bio-Rad) was added to the membranes and the Western Blot was resolved using the ChemiDoc (Bio-Rad).

### CRISPR/Cas9 screen

The appropriate number of cells to achieve 250-fold representation of the library, multiplied by five to account for 20% transduction efficiency, were transduced at approximately 40-60% confluence in the presence of polybrene (8 μg/mL) with the appropriate volume of the lentiviral-packaged sgRNA library. Cells were incubated overnight, followed by replacement of the lentivirus-containing medium with fresh medium containing puromycin (2-4 μg/mL). The lentivirus volume to achieve a MOI of 0.2, as well as the puromycin concentration to achieve a complete selection in 3 days was previously determined for each cell line. Transductions were performed in triplicate. After puromycin selection, cells were split into the indicated arms (for each arm, the appropriate number of cells to keep a 250-fold representation of the library was plated at approximately 10-20% confluence) and a T0 (reference) time point was harvested. Cells were maintained as indicated. In case a passage was required, cells were reseeded at the appropriate number to keep at least a 500-fold representation of the library. Cells (enough to keep at least a 500-fold representation of the library, to account for losses during DNA extraction) were collected when indicated, washed with PBS, pelleted and stored at –80°C until DNA extraction.

### DNA extraction, PCR amplification and Illumina sequencing

Genomic DNA (gDNA) was extracted (Zymo Research, D3024) from cell pellets according to the manufacturer’s instructions. For every sample, gDNA was quantified and the necessary DNA to maintain a 250-fold representation of the library was used for subsequent procedures (for this we assumed that each cell contains 6.6 pg genomic DNA). Each sample was divided over 50 μl PCR reactions (using a maximum of 1 µg gDNA per reaction) using barcoded forward primers to be able to deconvolute multiplexed samples after next generation sequencing (for primers and barcodes used, see Supplementary Table 3). PCR mixture per reaction: 10 μl 5x HF Buffer, 1 μl 10 μM forward primer, 1 μl 10 μM reverse primer, 0.5 μl Phusion polymerase (Thermo Fisher, F-530XL), 1 μl 10mM dNTPs, adding H2O and template to 50 μl. Cycling conditions: 30 sec at 98°C, 20× (30 sec at 98°C, 30 sec at 60°C, 1 min at 72°C), 5 min at 72 °C. The products of all reactions from the same sample were pooled and 2 μl of this pool was used in a subsequent PCR reaction using primers containing adapters for next generation sequencing (Supplementary Table 2). The same cycling protocol was used, this time for 15 cycles. Next, PCR products were purified using the ISOLATE II PCR and Gel Kit (Bioline, BIO-52060) according to the manufacturer’s instructions. DNA concentrations were measured and, based on this, samples were equimolarly pooled and subjected to Illumina next generation sequencing (HiSeq 2500 High Output Mode, Single-Read, 65 bp). Mapped read-counts were subsequently used as input for the further analyses.

### Bioinformatics Analysis

#### Screen Analysis

For each CRISPR screen the sgRNA count data for each sample was normalized for sequence depth using DESeq2, with the difference that the median instead of the total value of a sample was used. Then we took the results from the DESeq2 analysis and sorted it on the DESeq2 statistic in decreasing order putting the most enriched sgRNA at the top. We then ran the MAGeCK Robust Rank (RRA) tool on this list to test for enrichment of the sgRNAs of a gene towards the top for which RRA will generate a multiple testing corrected p-value (FDR).

#### TCGA data

Data generated by The Cancer Genome Atlas (TCGA) pilot project, established by the NCI and the National Human Genome Research Institute, were downloaded. Information about The TCGA and the investigators and institutions who constitute the TCGA research network can be found at https://cancergenome.nih.gov/. Level 3 pre-processed data were obtained from the TCGA Data Portal using the TCGAbiolinks R/Bioconductor package (Colaprico, Silva et al. 2016). The functions GDCquery, GDCdownload and GDCprepare were used to import data from the “TCGA-COAD” cohort into R (http://www.r-project.org) for further analysis.

#### MSI status

MSI status was derived from pre-calculated MSIsensor scores (Niu, Ye et al. 2014) found in (Bailey, Tokheim et al. 2018). MSI high (MSI-H) status was defined as MSIsensor score ≥ 10 and a MSI low (MSI-L) status was defined as MSIsensor score < 10 (Middha, Zhang et al. 2017).

#### Gene mutation status

Only non-synonymous mutations were considered. Tumour samples with a non-synonymous SNP in any of the following genes were considered as MMR mutant samples: MLH1, MLH3, PMS1, MS2, MSH2, MSH3, MSH6. Tumour samples with a non-synonymous SNP in MED12 were considered as MED12 mutant samples. Tumour samples with non-synonymous SNPs in both MED12 as well as an MMR gene were considered MED12+MMR double mutant samples.

#### Mutational signature analysis

A similar workflow for single bases substitution signature analysis was used as described elsewhere (Maura, Degasperi et al. 2019). In short, de novo signature extraction and assignment of the extracted signatures to the COSMIC reference catalog was employed using the mutationalPatterns R/Bioconductor package (Blokzijl, Janssen et al. 2018). Hereafter, the subset of COSMIC signatures identified from the extraction process was fitted using the deconstructSigs R/Bioconductor package (Rosenthal, McGranahan et al. 2016).

#### Gene set enrichment analysis

STAR pre-processed gene expression count data were downloaded from the TCGA Data Portal using the TCGAbiolinks R/Bioconductor package (Colaprico, Silva et al. 2016). Count normalization was performed with the DeSeq2 R/Bioconductor package (Love, Huber et al. 2014). GSEA was performed with GSEA software (version 4.2.2) (Subramanian, Tamayo et al. 2005). The Hallmark (version 7.5.1) and KEGG (version 7.5.1) gene sets were queried.

### Generation of custom MSI-focused sgRNA library

For the design of the custom sgRNA library we used the Broad GPP sgRNA design portal. The sgRNA sequences were ordered as a pool of oligonucleotides (Agilent) with flanking sequences to enable PCR amplification and Gibson assembly into pLentiGuide-Puro (pLG, addgene #52963). The pooled oligo library was amplified using pLG_U6_foward 5’-GGCTTTATATATCTTGTGGAAAGGACGAAACACCG-3’ and pLG-TRACR_Reverse 5’-GACTAGCCTTATTTTAACTTGCTATTTCTAGCTCTAAAAC-3’. The fragments were purified and cloned into pLG. The representation of the custom sgRNA library was validated by next generation sequencing.

### G23 MSI Activator vector generation

The G23 MSI Activator was assembled as described previously (31). In short, a pCDH plasmid containing Cre recombinase was digested with BamHI to split the Cre gene. The resultant linear plasmid was re-assembled with a synthetic intron through isothermal assembly. The Cre start codon was replaced by an EcoRI-spacer-XhoI site, allowing subsequent introduction of a synthetic “MSI tract” (repeat of 23 guanines) by annealed oligo cloning. The second and third ATG codons of the Cre recombinase gene were replaced by TGT codons (to prevent low level Cre translation occurring from alternative start sites). Together, this resulted in the following cassette: 5’ Kozak, an EcoRI restriction site, spacer, an XhoI restriction site, a Cre recombinase gene 3’. In its base configuration, the Cre recombinase gene is out of frame.

### MSI Reporter vector generation

LoxP sites were introduced into the multiple cloning sites of the pCDH-CMVp-MCS-PGK-BlastR vector, through two sequential rounds of annealed oligo cloning. First, the 3’ loxP site was introduced using EcoRI and BamHI restriction sites. The introduced region additionally included a short spacer sequence and a XhoI site directly upstream of the LoxP. Second, the 5’ loxP site was introduced using the XbaI and EcoRI restriction sites. This resulted in a modified MCS, comprising XbaI-loxP-EcoRI-spacer-XhoI-loxP-BamHI. Next, a scrambled non-sense open reading frame, terminated by 3 successive stop codons (TGA-TAA-TAG), was introduced into the EcoRI and XhoI sites. Then, a katushka open reading frame was introduced via the BamHI site. Finally, we PCRed a T2A-neomycin fragment using primers with homology to katushka and the PGK promoter and introduced it using Gibson assembly. This resulted in a vector containing from 5’ to 3’; The CMV promoter, a floxed non-sense open reading frame, a Katushka open reading frame, a T2A peptide, a neomycin resistance gene, the PGK promoter and a blasticidin resistance gene.

### MSI PCR test

MSI status was determined using the MSI Analysis kit (MD1641, Promega) according to the manufacturer’s instructions.

## Acknowledgements

This research was supported by an institutional grant of the Dutch Cancer Society and of the Dutch Ministry of Health, Welfare and Sport. Dr. Loredana Vecchione is participant in the BIH Charité (Junior) (Digital) Clinician Scientist Program funded by the Charité – Universitätsmedizin Berlin, and the Berlin Institute of Health at Charité (BIH)

## Supplementary Information

**Supplemental Figure 1:**
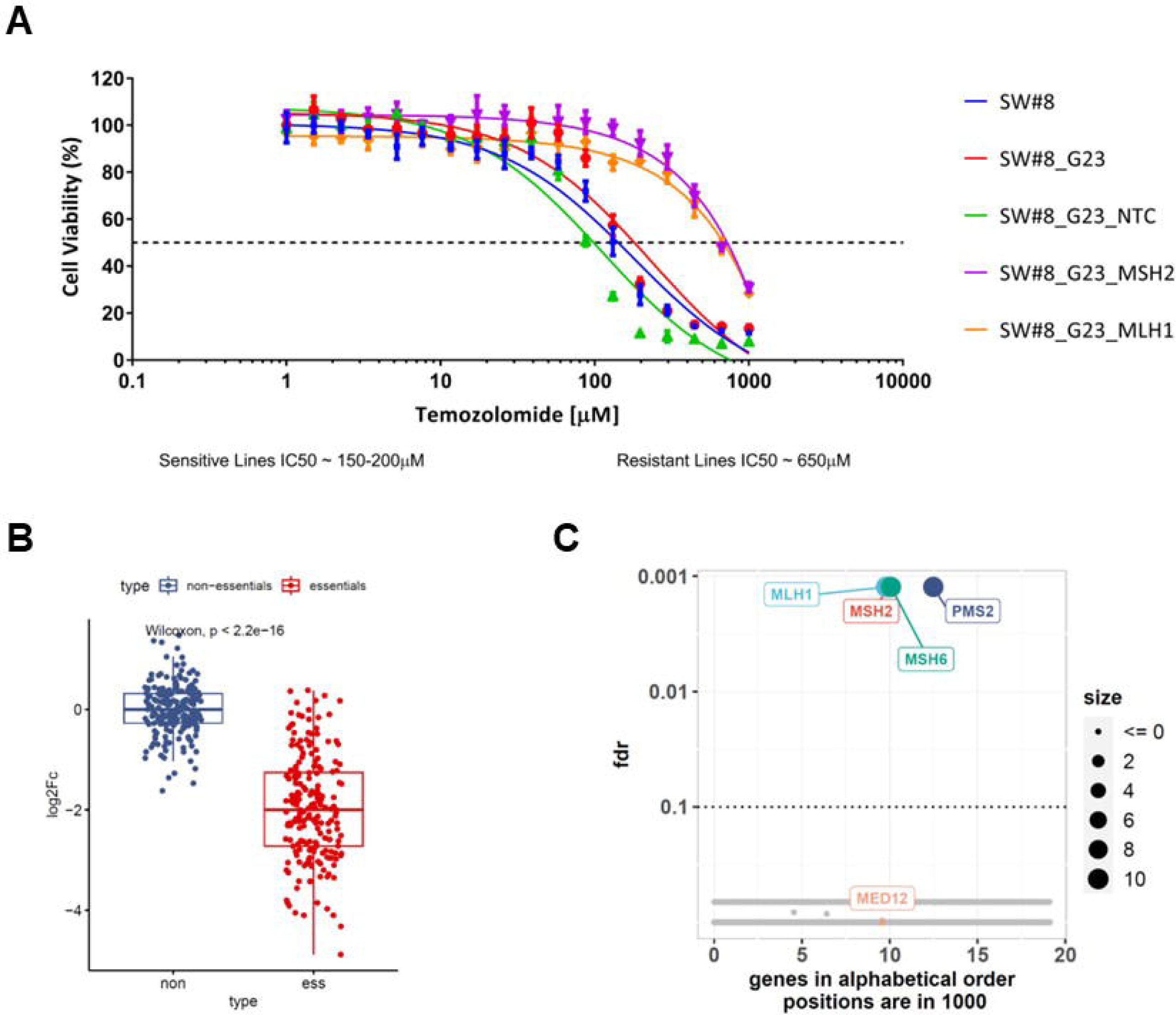
Using CRISPR screens to identify potential regulators of microsatellite instability. A. Cells were cultured with increasing concentrations of Temozolomide for 4 days, after which cell viability was measured using CellTiter-Blue®. Standard deviation (SD) from 3 replicates is plotted. B,C. Analysis of the genome-wide screen. Cas9 expressing SW#8 cells were screened with the Brunello whole-genome sgRNA library. In (B) the analysis of the depletion (log2 Fold Change) of the sgRNAs targeting essential genes over non-essential genes is displayed. Box plot shows the median (horizontal line), interquartile range (hinges), and the smallest and largest values no more than 1.5 times the interquartile range (whiskers). Comparisons were made using the Wilcoxon test. In (C) Robust rank results for enrichment of each gene in the Temozolomide treated arm (T48) compared to the reference (T12) is displayed.

**Supplemental Figure 2:**
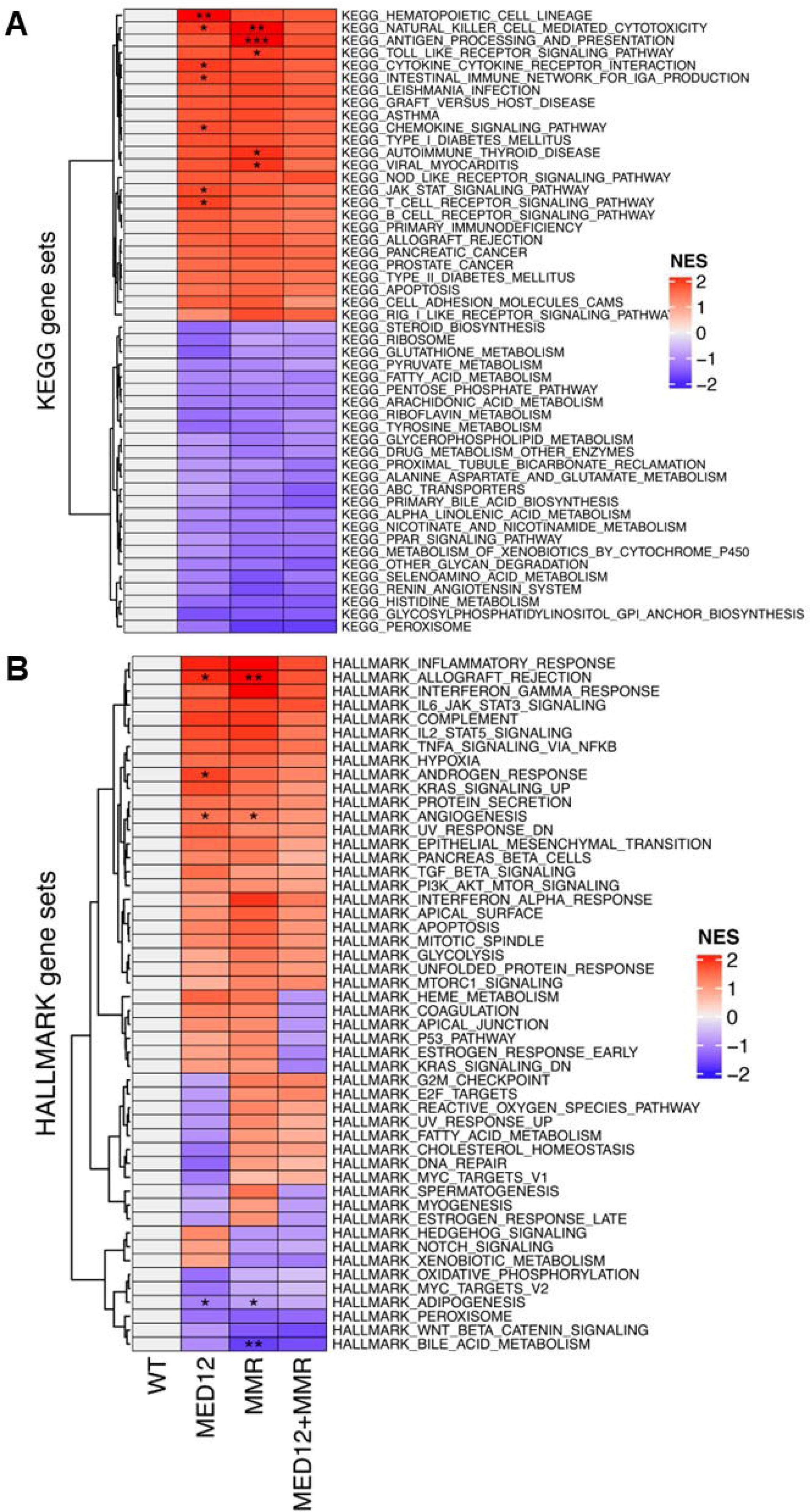
MED12 mutant colon adenocarcinomas contain an inflammatory gene expression signature. Gene set enrichment analysis (GSEA) relative to wild type samples (i.e. no mutation in MED12 and MMR genes). A. The normalized enrichment scores (NES) are depicted in a heatmap that is filtered for the top 25 over– and under-represented KEGG pathways. B. The normalized enrichment scores of hallmark gene sets are depicted in a heatmap. (*FDR ≤ 0.05, **FDR ≤ 0.01, ***FDR ≤ 0.001).

**Table S1:** Brunello screen analysis, temozolomide-treated arm.

**Table S2:** Brunello screen analysis, untreated arm.

**Table S3:** MSI-focused library.

**Table S4:** MSI-focused screen analysis.

